# SARM1 orthosteric base exchange inhibitors cause subinhibitory SARM1 activation

**DOI:** 10.1101/2024.08.06.606489

**Authors:** Rebecca Leahey, Martin Weber, Chang Hoon Cho, Seong Hur, Amber Cramer, Karla Manzanares, Brett Babin, Gladys Boenig, Taylor Kring, Liling Liu, Yusi Cui, Anjani Ganti, John P. Evans, Marika Nespi, Justin Ly, Alicia A. Nugent, Samantha Green, Bryan Chan, Casper C. Hoogenraad, Anton Delwig, Flora I. Hinz

## Abstract

SARM1, an octameric NADase, is a key regulator of axon degeneration and an emerging target in small molecule drug discovery to treat a wide range of neurodegenerative diseases. Recently, a structurally diverse series of adduct-forming, orthosteric SARM1 inhibitors have been discovered. Here, we show that subinhibitory concentrations of these orthosteric inhibitors, under mildly SARM1 activating conditions, cause sustained SARM1 activation. This synergistic adverse effect leads to increased nicotinamide adenine dinucleotide (NAD) consumption, neurodegeneration and release of the biomarker neurofilament-light (NfL) in cultured cortical neurons. In two distinct animal models, we found that low-dose treatment with these orthosteric SARM1 inhibitors results in increased plasma NfL and adverse events when combined with cellular stress or injury conditions. This may present a critical liability for orthosteric SARM1 inhibitors in certain patient populations.

## MAIN

Axon degeneration, a key, early component of most neurodegenerative diseases, is an active process regulated by the NADase SARM1 (Essuman et al., 2017). SARM1 forms a homo-octamer (Sporny et al., 2019) and acts as a metabolic sensor. In healthy neurons under metabolic steady-state conditions, SARM1 is autoinhibited by NAD binding to an allosteric site on the ARM domain. Under conditions of cellular stress, NAD levels drop and nicotinamide mononucleotide (NMN) displaces NAD at this site to cause conformational rearrangement of the octamer, dimerization of the catalytic TIR domains and formation of six active sites (Figure 1a, Figley et al., 2021, Sporny et al., 2020). Upon activation, rapid hydrolysis of NAD by SARM1 leads to mitochondrial dysfunction, shutdown of ATP synthesis, Calcium influx (Ko et al., 2021), dismantling of the cytoskeleton and subsequent release of NfL, a widely accepted biomarker of axon degeneration (Khalil et al., 2024; Figure 1a).

**Figure 1.**
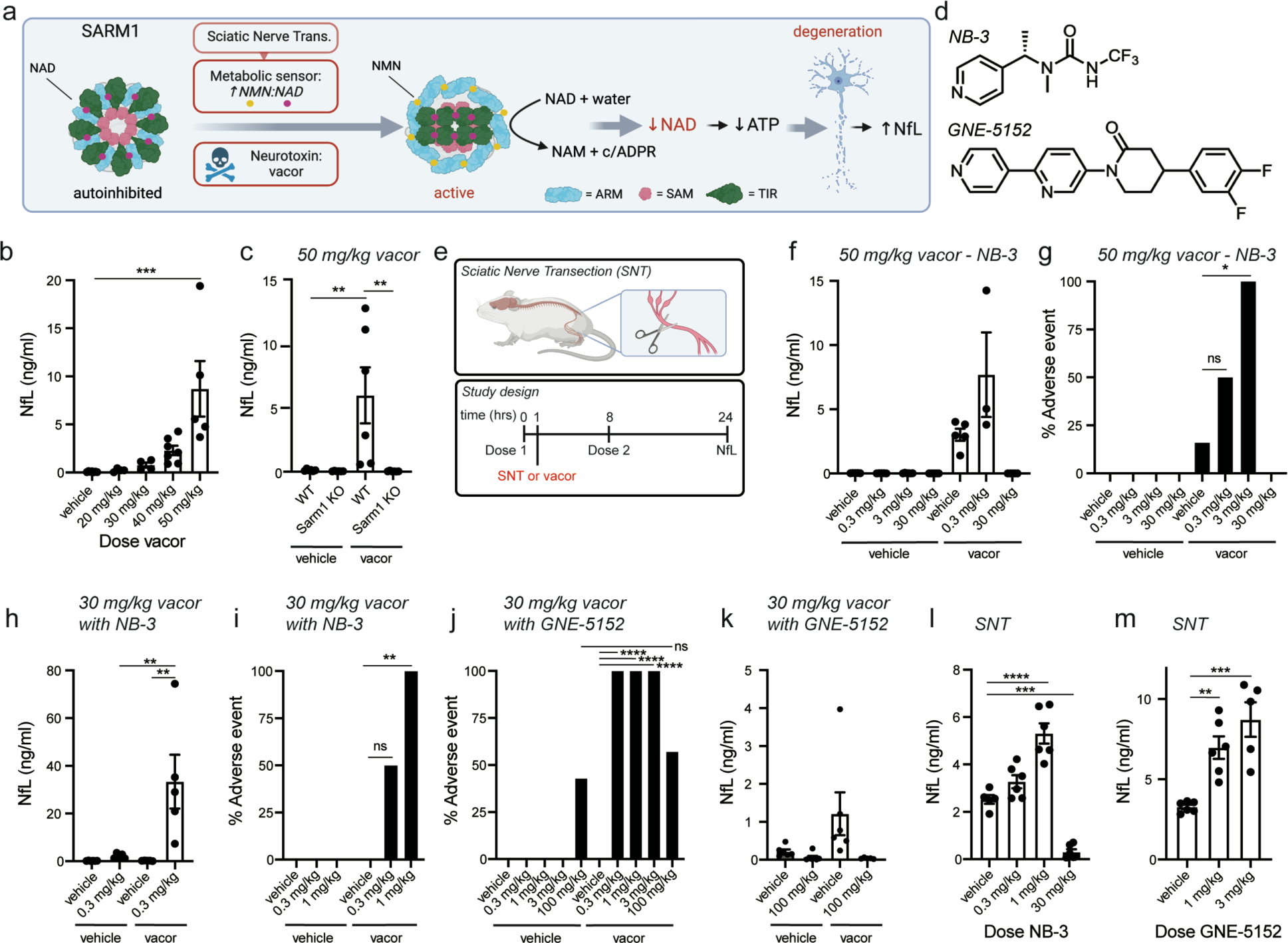
Low concentrations of SARM1 BE inhibitors increase adverse events and plasma NfL in mouse models of acute neuronal injury. (a) Schematic of SARM1 activation and downstream consequences. (b) Plasma NfL 24hrs after oral dosing of vacor (n=4-7 mice/group). Dunnett’s multiple comparisons test relative to vehicle treatment, ***p<0.001. (c) Plasma NfL 24hrs after oral dosing of WT or Sarm1 KO mice with 50 mg/kg vacor (n=6 mice/group). Tukey’s multiple comparisons test **p<0.01. (d) Structures of SARM1 BEI inhibitors. (e) Schematic of SNT (top) and subsequent study design time course (bottom). Plasma NfL (f) and percent adverse events observed (g) in mice co-dosed with 0-30 mg/kg NB-3 and 0 or 50 mg/kg vacor (f, n=3-6 mice/group; g, n=6 mice/group; Fisher’s Exact Test). Plasma NfL (h) and percent adverse events observed (i) in mice co-dosed with 0-1 mg/kg NB-3 and 0 or 30 mg/kg vacor (h, n=5-6 mice/group, Tukey’s multiple comparisons test **p<0.01; i, n=6 mice/group, Fisher’s Exact Test). Percent adverse events observed (j) and plasma NfL (k) in mice co-dosed with 0-100 mg/kg GNE-5152 and 30 mg/kg vacor (j, n=6-13 mice/group, Fisher’s Exact Test; k, n=5-7 mice/group). (l) Plasma NfL after SNT in mice dosed with 0-30 mg/kg NB-3 (n=5-6 mice/group). Tukey’s multiple comparisons test ***p=0.002 and ****p<0.0001. (m) Plasma NfL after SNT in mice dosed with 0-3 mg/kg GNE-5152 (n=5-6 mice/group). Tukey’s multiple comparisons test **p=0.0046 and ***p=0.0003.

SARM1 loss or inhibition has been shown to protect axons and reduce plasma NfL levels after nerve injury, in response to focal application of neurotoxins and models of neurodegeneration (Wu et al., 2021, Loreto et al., 2021, Bratkowski et al., 2022). Due to its central role in regulating axon degeneration, SARM1 is an emerging target in small molecule drug discovery for treating a variety of neurodegenerative diseases. Based on a survey of disclosed structures, one common inhibition approach is a mechanism-based inactivation strategy (Hughes et al., 2019; Kozak et al., 2022), which takes advantage of the base-exchange (BE) activity of SARM1. This class of inhibitors consists of prodrugs that are converted by SARM1 into nicotinamide-substituted NAD analogs, which are potent active site (orthosteric) SARM1 inhibitors (Shi et al., 2022; Bratkowski et al., 2022).

To study the inhibitory effects of these SARM1 BE inhibitors (BEIs) we developed a mouse model of systemic vacor dosing to activate SARM1. Vacor mononucleotide, a metabolite of the neurotoxin vacor, has recently been shown to activate SARM1 by mimicking NMN (Loreto et al., 2021). 50 mg/kg vacor administered once orally was generally tolerated and led to significant increases in plasma NfL levels at 24hrs (Figure 1b). This NfL increase was SARM1-dependent (Figure 1c). To investigate the dose-response effects of SARM1 BEIs in this model, we dosed NB-3, a previously described BEI (Bratkowski et al., 2022, Figure 1d), twice orally at 0 and 8hrs in combination with 50 mg/kg vacor at 1hr and collected plasma to measure NfL at 24hrs (Figure 1e, Supplementary Figure 1a). While 30 mg/kg NB-3 significantly reduced plasma NfL, indicative of neuroprotection, lower doses of NB-3 caused not only increased NfL release but also incidents of serious adverse events, such as lethargy, immobility, and death (Figure 1f,g). In the absence of vacor co-dosing, NB-3 did not elevate plasma NfL or cause adverse events (Figure 1g).

To further investigate this phenomenon, we dosed NB-3 and GNE-5152, a structurally distinct BEI developed internally (Figure 1d, Supplementary Figure 1b), in two additional mouse models of SARM1 activation: a low-dose vacor and sciatic nerve transection (SNT) model. In the first model, the dose of vacor was reduced to 30 mg/kg to increase the dynamic range of NfL measurements and further decrease the likelihood of vacor-induced adverse events. In this model, subinhibitory doses of NB-3 increased plasma NfL and subinhibitory doses of both NB-3 and GNE-5152 caused serious adverse events (Figure 1h-j). While 100 mg/kg GNE-5152 resulted in decreased plasma NfL when co-dosed with vacor (Figure 1k), this high dose of GNE-5152 caused adverse events independent of vacor co-dosing. Similar BEI-induced plasma NfL elevation was also observed in the SNT model. While high doses of NB-3 reduced NfL after SNT, indicating protection of the injured nerve, lower doses of both NB-3 and GNE-5152 significantly elevated plasma NfL (Figure 1l,m). Together, these results demonstrate that subinhibitory concentrations of orthosteric SARM1 BEIs in the presence of SARM1 activating stimuli result in increased plasma NfL and serious adverse events *in vivo*.

To elucidate the mechanism behind this synergistic adverse effect, we evaluated SARM1 BEIs in biochemical and cellular assays designed to measure SARM1 activity and SARM1-dependent cell death. The enzymatic activity of recombinant human SARM1 was assessed by continuously monitoring the conversion of the synthetic fluorogenic SARM1 substrate, PC6, into its fluorescent product, PAD6 (Li et al., 2021), and by quantifying the production of nicotinamide mononucleotide (NAM) resulting from the SARM1-catalyzed conversion of NAD. Under physiological NAD concentrations (200 µM) and submaximal SARM1 activation (24 µM NMN), subinhibitory concentrations of GNE-5152 and NB-3 led to sustained SARM1 activation (Figure 2a,b). To verify that this sustained SARM1 activity as indicated by PAD6 formation corresponds to increased NAD consumption, we quantified NAM production 800 minutes after initiating the biochemical reaction. NAM levels were higher in the presence of low concentrations of BEIs (Figure 2c,d), confirming increased SARM1 activity under these conditions. The BEI-induced increase of SARM1 activity was also observed in cellular viability assays under conditions of partial SARM1 activation. SARM1 can be activated by vacor in a dose-dependent manner, leading to reduced cell viability at 24hrs (Supplementary Figure 1c,d). When SARM1 was submaximally activated by vacor, high concentrations of GNE-5152 and NB-3 inhibited SARM1, thereby rescuing cell viability. In contrast, subinhibitory concentrations of BEIs potentiated SARM1-induced cell death (Figure 2e,f). To determine whether this activation effect is consistent across different compounds, we screened a small library of structurally diverse, internally developed BEIs. All tested BEI compounds showed an activation effect at subinhibitory concentrations (Figure 2g).

**Figure 2.**
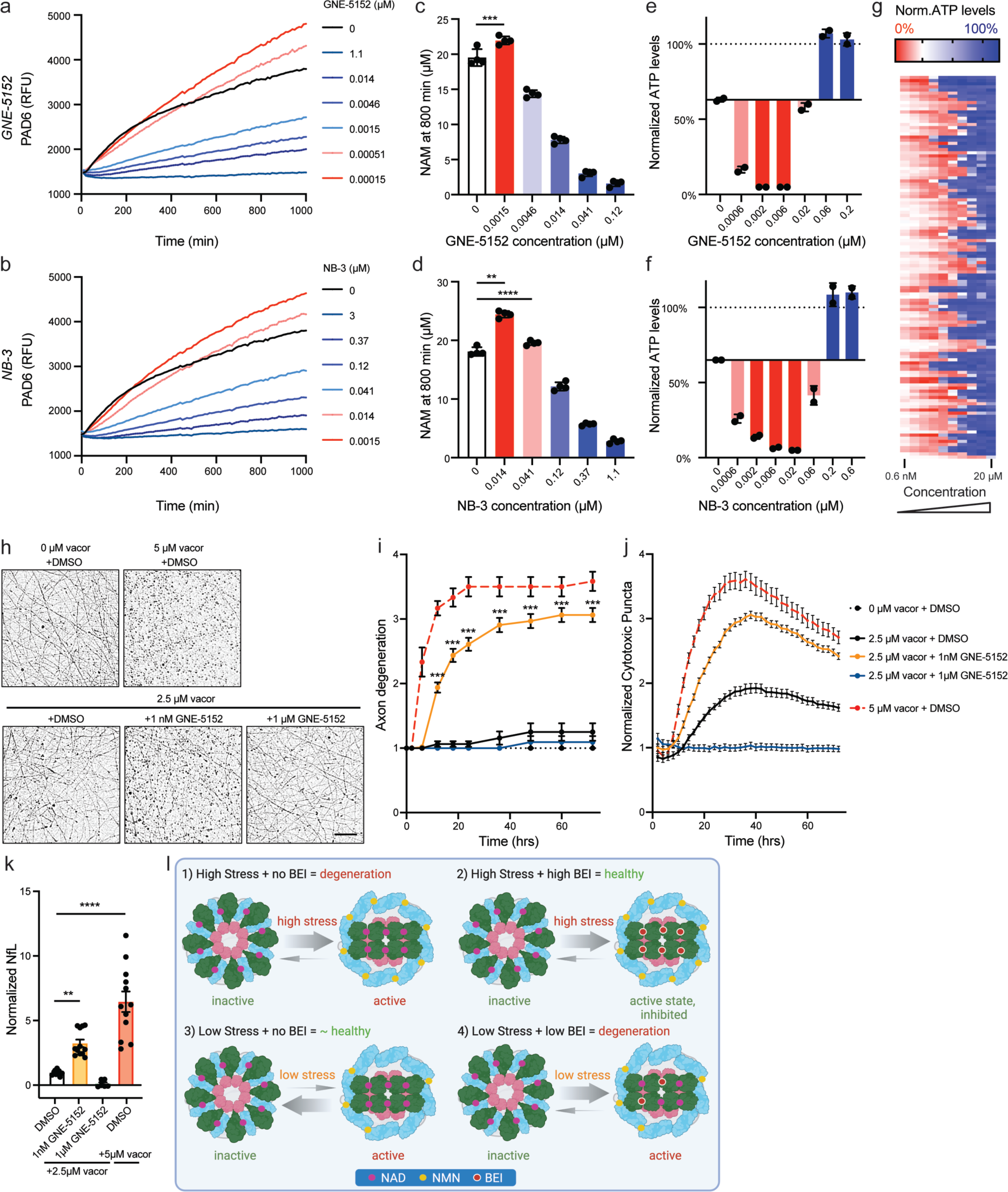
SARM1 BE inhibitors paradoxically activate SARM1 at subinhibitory concentrations in biochemical and cellular assays causing cell death and NfL release. Quantification of PAD6 fluorescence after incubation of recombinant hSARM1 with varying concentrations of either (a) GNE-5152 or (b) NB-3. n=4. NAM concentration at 800 minutes after incubation of recombinant hSARM1 with varying concentrations of either (c) GNE-5152 or (d) NB-3. n=4; one-way ANOVA, ****p<0.0001, ***p<0.001, **p<0.01. Normalized ATP levels 24hrs after treatment of SH-SY5Y cells with 10 µM vacor and varying concentrations of either (e) GNE-5152 or (f) NB-3. n=2. (g) Heat map of mean normalized ATP levels 24hrs after treatment of SK-N-SH cells with 5 µM vacor and varying concentrations of 122 representative BE compounds (each row represents 1 compound, n=2). Sample images (h) and quantification (i) of axon degeneration of WT mouse cortical neurons treated with 0-5 µM vacor in the presence of 0-1 µM GNE-5152. n=12-64 images/condition, Mann-Whitney test ***p<0.001 comparing 2.5 µM vacor + DMSO to 2.5 µM vacor + 1 nM GNE-5152. Scale bar = 50 µM. (j) Quantification of cytotoxic puncta of WT mouse cortical neurons treated with 2.5-5 µM vacor in the presence of 0-1 µM GNE-5152, normalized to 0 µM vacor. n=14-20 images/condition; for all time points after 8 hrs, p<0.0001 when comparing 2.5 µM vacor + DMSO to 2.5 µM vacor + 1 nM GNE-5152. (k) NfL release from mouse cortical cultures 48hrs after treatment, normalized to 2.5 µM vacor + DMSO. n=6-12 wells/condition; Dunnett’s multiple comparisons test, **p<0.01, ****p<0.0001. (l) Schematic of SARM1 BEI stabilizing active SARM1 under low stress conditions.

Next, we investigated whether SARM1 BEIs cause deleterious effects in primary neurons, a therapeutically relevant cell type. Using previously established methods to monitor axon degeneration (Hinz et al., 2024), we found that subinhibitory concentrations of GNE-5152 exacerbated axon degeneration only under partially SARM1 activating conditions (2.5 µM vacor) (Figure 2h,i; Supplementary Figure 2a-c). In the absence of stress, concentrations between 1nM-10 µM GNE-5152 did not induce axon degeneration (Supplementary Figure 2a). Under high SARM1 activation conditions (5 µM vacor), subinhibitory concentrations of GNE-5152 did not further exacerbate axon degeneration consistent with a ceiling effect, while high inhibitor concentrations prevented axon degeneration (Supplementary Figure 2c). Similar effects were observed with SARM1 BEI NB-3 (Supplementary Figure 2d-f), again confirming that this interaction effect between subinhibitory concentrations of SARM1 BEI and stress state is not specific to one chemical series. Furthermore, subinhibitory concentrations of SARM1 BEIs GNE-5152 and NB-3 in the presence of mild stress resulted in increased cytotoxic puncta, a measure of cell body death (Figure 2j, Supplementary Figure 2g) and increased NfL release, as compared to mild stress conditions alone (Figure 2k, Supplementary Figure 2h).

This interaction effect between mild stress and subinhibitory SARM1 BEI concentrations was SARM1-dependent (Supplementary Figure 3) and mediated by early loss of NAD levels (Supplementary Figure 4). Furthermore, the interaction effect leading to increased axon degeneration was not dependent on SARM1 activation by vacor specifically, but appeared to be general for other SARM1 activating stress conditions. Cortical neurons treated with low concentrations (2.5 nM) of vincristine, a microtubule destabilizing chemotherapy agent known to cause peripheral neuropathy in patients and SARM1-dependent axon degeneration *in vitro*, and subinhibitory concentrations of GNE-5152 showed exacerbated axon degeneration as compared to 2.5 nM vincristine treatment in the presence of DMSO (Supplementary Figure 5). This effect was not limited to rodent neurons. When iPSC-derived human cortical neurons were treated with mildly activating concentrations of vacor (25 µM), increased axon degeneration was observed when neurons were co-treated with low concentrations (1 and 2 nM) of SARM1 BEI (Supplementary Figure 6).

In summary, we find that BEIs cause paradoxical and sustained activation of SARM1. At concentrations lower than effective inhibitory concentration, this BEI-induced SARM1 activity leads to increased cell death, NfL release and severe adverse events *in vivo*. While the exact mechanism of the observed sustained SARM1 activation is unclear, we speculate that sub-stoichiometric binding of BEIs stabilize the active form of SARM1 while not blocking all catalytic sites. This allows the unoccupied catalytic sites to remain active and continue to consume NAD (Figure 2l). Similar ‘paradoxical’ activation by substoichiometric inhibition has been observed for a variety of other multimeric enzymes (Merdanovic et al., 2020, Hatzivassiliouet al., 2010).

In the case of SARM1 BEIs, this subinhibitory activation effect is dependent on partial SARM1 activation and is not detected in the absence of cellular stress or in healthy animals. However, under standard inhibitor screening conditions optimized to cause maximal enzyme activity to increase the assay window, any additional SARM1 activity caused by BEIs may not be detected. Additionally, biochemical and cellular assays need to be of sufficient duration to measure the delayed phase of sustained SARM1 activity, which can be easily missed if the assays are of short duration. In animal models, NfL measurements are often used as a proxy for neuronal degeneration. However, NfL release is frequently transient in these models (Sasaki et al., 2020) and if NfL measurements are confined to peak NfL release time points, left shifts of NfL release curves might be interpreted as NfL release reduction at peak. Therefore, this subinhibitory activation effect may easily be missed in assays and models designed to detect SARM1 inhibition during early preclinical development.

In our studies, all tested BEIs increased SARM1 activity at subinhibitory concentrations, leading to serious adverse events in mice. While there is currently no direct evidence whether and to what extent SARM1 is activated in patients with neurodegenerative diseases, the assumption that SARM1 activity is driving axon degeneration in this patient population underlies the therapeutic hypothesis for SARM1 inhibition as a treatment for neurodegeneration. Therefore, our observations that subinhibitory concentrations of orthosteric SARM1 BEIs cause sustained SARM1 activity and increase plasma NfL are potentially concerning. Critically, as this interaction effect requires low levels of SARM1 activation, studies that assess the safety of BEIs in healthy controls are unlikely to ascertain the translatability of these findings to patients with ongoing neurodegeneration.

## ONLINE METHODS

### Reagents

Vincristine (V8879), NAD (10127965001), and NMN were purchased from Sigma-Aldrich. Vacor was synthesized as described in Hinz et al., 2024. NB-3 was prepared according to Bratkowski et al., 2022.

Synthesis of GNE-5152:

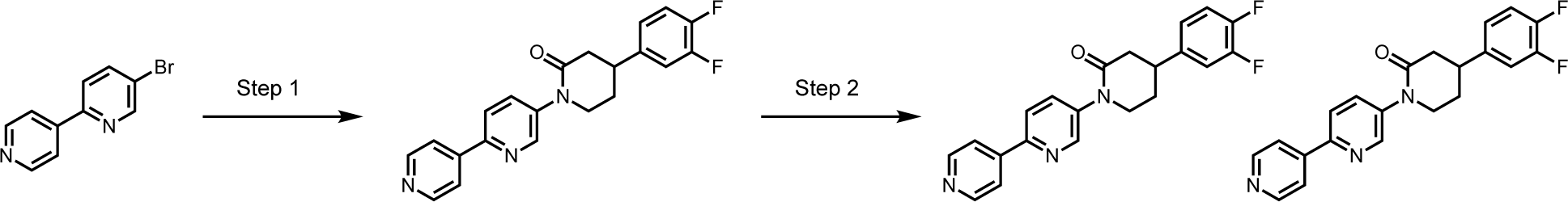

#### Step 1: Synthesis of 1-([2,4’-bipyridin]-5-yl)-4-(3,4-difluorophenyl)piperidin-2-one

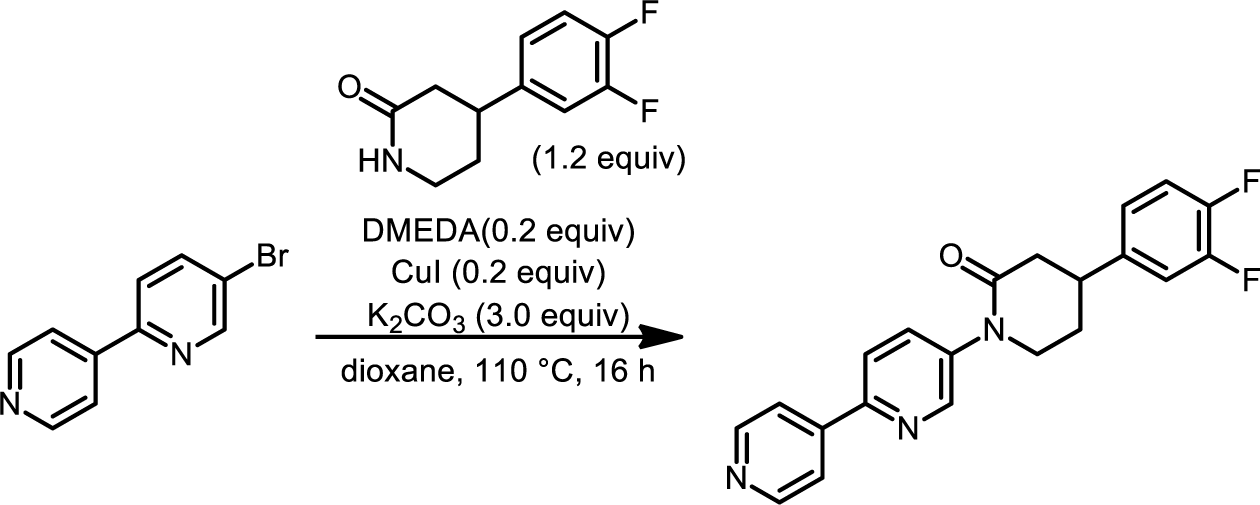

To a mixture of 4-(3,4-difluorophenyl)piperidin-2-one (216 mg, 1.02 mmol), 5-bromo-2,4’-bipyridine (200 mg, 0.85 mmol) in 1,4-dioxane (4 mL) was added DMEDA (0.02 mL, 0.2 mmol), K_2_CO_3_ (353 mg, 2.6 mmol) and CuI (32 mg, 0.2 mmol). The mixture was stirred at 110 °C for 16 h under nitrogen atmosphere. After cooling to room temperature, the reaction mixture was filtered and the filtrate was concentrated under reduced pressure. The residue was purified by flash column chromatography (SiO_2_, 0-36% EE (25% ethanol in ethyl acetate) in petroleum ether) to afford 1-([2,4’-bipyridin]-5-yl)-4-(3,4-difluorophenyl)piperidin-2-one (200 mg, 64% yield) as a white solid.

#### Step 2: Chiral Separation of 1-([2,4’-bipyridin]-5-yl)-4-(3,4-difluorophenyl)piperidin-2-one

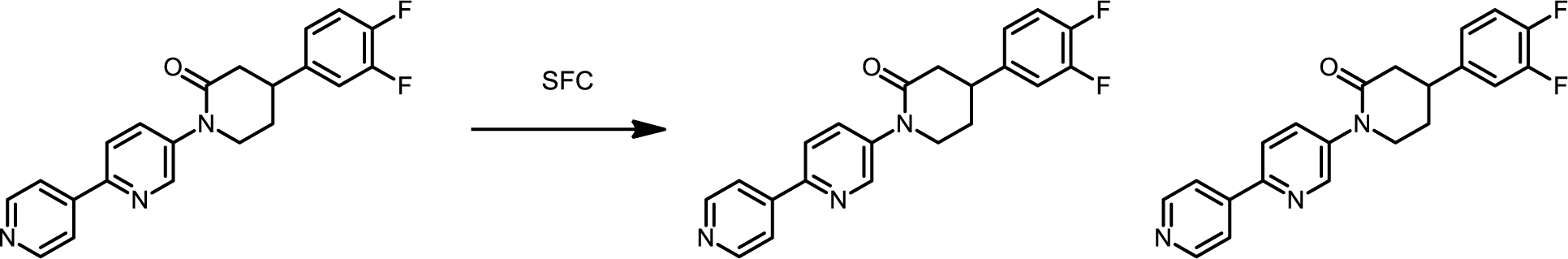

1-([2,4’-bipyridin]-5-yl)-4-(3,4-difluorophenyl)piperidin-2-one (200 mg, 0.55 mmol) was separated by chiral SFC (Cellulose-2 (250 mm x 30 mm, 10 um), Supercritical CO_2_ / EtOH + 0.1% NH_4_OH = 50/50; 150 mL / min) to afford 1-([2,4’-bipyridin]-5-yl)-4-(3,4-difluorophenyl)piperidin-2-one (**GNE-5151**,peak 1, Rt = 3.622 min, 57.9 mg, 29% yield) and 1-([2,4’-bipyridin]-5-yl)-4-(3,4-difluorophenyl)piperidin-2-one (**GNE-5152**, peak 2, Rt = 4.371 min, 64.8 mg, 32%) as a white solid.

**GNE-5151 ^1^H NMR** (DMSO-*d*_6_, 400 MHz): *δ* 8.75 (d, *J* = 2.4 Hz, 1H), 8.71 (d, *J* = 4.8 Hz, 2H), 8.18 (d, *J* = 8.4 Hz, 1H), 8.06 (d, *J* = 4.8 Hz, 2H), 7.99 - 7.91 (m, 1H), 7.51 - 7.38 (m, 2H), 7.22 - 7.17 (m, 1H), 4.00 - 3.88 (m, 1H), 3.79 - 3.67 (m, 1H), 3.40 - 3.36 (m, 1H), 2.74 - 2.65 (m, 2H), 2.20 - 2.10 (m, 2H). **LCMS**: (ESI, m/z) [M+H]^+^ = 366.1

**GNE-5152: ^1^H NMR** (DMSO-*d*_6_, 400 MHz): *δ* 8.75 (d, *J* = 2.4 Hz, 1H), 8.70 (d, *J* = 5.6 Hz, 2H), 8.18 (d, *J* = 8.4 Hz, 1H), 8.06 (d, *J* = 6.0 Hz, 2H), 7.99 - 7.91 (m, 1H), 7.51 - 7.36 (m, 2H), 7.22 - 7.17 (m, 1H), 4.00 - 3.88 (m, 1H), 3.79 - 3.67 (m, 1H), 3.40 - 3.36 (m, 1H), 2.74 - 2.65 (m, 2H), 2.20 - 2.10 (m, 2H). **LCMS**: (ESI, m/z) [M+H]^+^ = 366.1

### Biochemical assays

The activity of recombinant human SARM1 was monitored using PC6 fluorogenic probe (Li et al., 2021) and mass spectrometry. Reaction mixtures contained 1 nM SARM1, 200 µM NAD (Roche, 10127973001), and 24 µM NMN (USBiological, 018099). 10 µM PC6 (synthesized as described in Li et al., 2021) was added for the kinetic fluorescence assay. The assay buffer was 50 mM Tris HCl pH 7.5 (Teknova, T1074), 0.5 mM DTT (Roche, 03117014007), 0.0036% Tween-20 (Sigma, P1379), 0.05% bovine gamma globulin (BGG, Sigma, P1379). Timecourse of PAD6 fluorescence (excitation/emission: 390/520 nm) was measured at room temperature for 16.5 hours at 5-minute intervals using Infinite M1000 plate reader (Tecan). Compound dilution series were prepared from 10 mM DMSO stock solutions and dispensed into assay plates using Echo555 (Labcyte, Beckman Coulter) at 1% final DMSO concentration. For mass spectrometry analysis, samples were quenched by addition of 0.1% formic acid in water with 10 mM ammonium formate. 4 µM nicotinamide-D4 (Cambridge Isotope Laboratories) was added as a loading standard. NAM was measured and quantified by a RapidFire (Agilent) system coupled to an AB Sciex TRIPLE QUAD 5500 mass spectrometer. Analyst 1.6.3 software packages (Applied Biosystems) and RapidFire 5.1 Acquisition and Integrator were used to control the RF-MS/MS system, as well as for data acquisition and processing. RapidFire conditions: graphite cartridge, pump 1 (Load/Wash): water with 10 mM ammonium formate, pump 3 (Elute): 50% acetonitrile in water with 10 mM ammonium formate. NAM (123/80) and NAM-D4 (127/84) ions were monitored. NAM measurements were normalized by NAM-D4 and then converted to NAM in μM based on an NAM calibration curve. To account for interference from NAD, NAM calibration samples contained both NAM and NAD at the final total concentration of 200 μM.

### Preparation of recombinant human SARM1 protein

pRK5 plasmid containing human SARM1 (E26-T274, EC 3.2.2.6) with N-terminal His and C-terminal Avi and Flag tags was transfected intExpi293F^TM^ cells (ThermoFisher Scientific #A14527) were cultured in Expi293 expression medium at 37 °C, 8% CO2. Cells were seeded at 2.5–3 × 10^6^ viable cells per ml and transfected with 0.8 mg/ml DNA construct using 1:3 PEI Max transfection reagent (PolySciences #24765). Post transfection, cells were fed and 4mM valproic acid was added. The cells were harvested after 48 h, by centrifuging at 500 × *g*, 15 min, 4 °C.

For purification, cell paste was resuspended in (100 mL per liter of cell paste) ice-cold lysis buffer (50 mM Tris-HCl pH 8.0, 200mM NaCl, 5 % Glycerol, 1.0 mM TCEP (*Tris-HCl*(2-carboxyethyl)phosphine), 10 Roche Complete EDTA-Free Protease Inhibitor Cocktail Tablets (Millipore Sigma #4693132001) and cells were homogenized then disrupted by double passage using Microfluidizer â Processor (Microfluidics Model M-110Y). Insoluble matter was separated by ultracentrifugation at 40,000 RPM for 1 hour. The soluble supernatant was decanted and passed through a monoclonal ANTI-FLAG^®^ M2 antibody (Millipore Sigma #F1804). The resin column was equilibrated with Lysis Buffer and the bound protein was eluted by 100 µg/mL of 3X FLAG^®^ Peptide (Sigma-Aldrich #F4799). The eluent protein was injected on a Superdex 200 16/60 (Amersham Biosciences # GE28-9893-35) column equilibrated in 25 mM Tris pH 8.0, 1 mM TCEP, 150 mM NaCl, 5% Glycerol.

### Neuroblastoma viability assay

To activate SARM1, SK-N-SH or SH-SY5Y cells were treated with the dose of vacor that reduced viability by 30-40% at 24 hours as assayed by the CellTiter-Glo reagent according to the manufacturer’s protocol (Promega, G7573). The dose of vacor was experimentally determined before each experiment. In this study, SARM1 in SH-SY5Y was activated with 10 µM vacor and with 5 µM in SK-N-SH. Compound dilution series were prepared from 10 mM DMSO stock solutions and dispensed into assay plates using Echo555 (Labcyte, Beckman Coulter) at 0.2% final DMSO concentration. Raw CellTiter-Glo luminesce values were normalized to no treatment controls (no vacor, 0.2% DMSO) and plotted as percent cell viability.

### Dissociated primary mouse cortical neuron cultures

To prepare primary embryonic cortical cultures, embryos were obtained from timed pregnant C57BL/6N mice from Charles River Laboratories (MA, USA) as well as timed pregnant *Sarm1*^-/-^ and *Sarm1*^+/+^ mice obtained from Jackson labs (B6.129X1-*Sarm1^tm1Aidi^*/J; Kim et al., 2007). Mice were housed on a regular light/dark cycle (14:10hr) with ad libitum access to food (LabDiet 5010) and water. All animal care/handling procedures were reviewed and approved by Genentech’s Institutional Animal Care and Use Committee, and were conducted in full compliance with regulatory statutes, Institutional Animal Care and Use Committee policies, and National Institutes of Health guidelines.

Cortices from day 15 embryos (E15) were dissected, stripped of meninges, washed 3x with cold HBSS (Invitrogen, 14170-112), and incubated for 10 min at 37°C in HBSS supplemented with 0.25% trypsin (Invitrogen, 15090-046) and DNase I (Roche, 104159). Tissue was washed 3x with HBSS and triturated in plating media containing DNase I (Neurobasal Medium (Thermo Fisher, 21103-049), 20% heat-inactivated horse serum (Thermo Fisher, 26050-088), 25 mM sucrose, and 0.25% Gibco GlutaMAX (Thermo Fisher, 35050-061)). Dissociated cells were centrifuged at 125 g for 5 min at 4°C, resuspended in plating medium, and plated in poly-L-lysine-coated (Millipore Sigma, P1274) plates. After 24 hrs, the plating medium was replaced with NBActiv4 (Brain Bits, NB4500). Cells were maintained at 37°C with 5% CO_2_ and the medium was renewed using 50% exchange every 3-4 days.

For ‘spot cultures’ used for axon degeneration assays, approximately 40,000 cells in a volume of 1.25 µL were plated on the center of a 48 well plate and incubated at 37°C with 5% CO_2_ for 20 minutes, after which the well was flooded with plating media. For cytotoxicity dye uptake and biochemical assays, cells were plated in 96 well plates at a density of 30,000 cells per well. All experiments were performed between DIV7-10.

### Human iPSC-derived cortical cultures

hiPSCs were cultured and maintained as previously described in Shan et al., 2024. Briefly, iP11N iPSCs were cultured and maintained in mTeSR™ Plus media (100-0276; Stemcell technologies) until reaching a confluency of ∼75%, and were then dissociated and replated at day 0 at a density of 40,000 cells / cm^2^ on iMatrix-511 (T304; Takara) coated plates in mTesR Plus supplemented with 3 µg/mL doxycycline and Y-27632 (S1049; Selleck Chemicals). On day 1, cells were switched into induction media containing SB431542, Noggin, XAV939, DATP and doxycycline. Induction media was changed daily for 5 days. On day 5, cells were switched into maturation media supplemented with 2 µg/ml doxycycline, 5 µM Cytosine β-D-arabinofuranoside hydrochloride (Ara-C: C6645-100MG; Sigma-Aldrich), and DAPT to promote neuronal differentiation and discourage neural progenitor expansion. The cells were cryopreserved on day 7 in CryoStor CS10 media (100-1061; Stemcell Technologies), creating D7-iNGN2 neuron banks. Spot cultures to measure axon degeneration were prepared as previously described in Hinz et al., 2024. Briefly, the D7-iNGN2 cells were thawed and seeded in droplets containing 50,000 cells in the middle of each poly-D-lysine (P7280; Sigma-Aldrich) and iMatrix coated well. Cells were incubated for 20 minutes to allow for attachment before the well was flooded with additional maturation media containing 2 µg/ml doxycycline. ⅓ of the media was replaced with fresh maturation media supplemented with doxycycline three times a week until day 14, after which cells were fed on the same schedule with maturation media without doxycycline until day 28.

### Axon degeneration assay

Axon stress treatment and SARM1 inhibitor addition was performed as part of a 50% media change. Immediately after treatment or injury, plates were placed into an S3 Incucyte Live-Cell Analysis System and phase contrast images were automatically captured every 2hrs for 72-96hrs. Two images per well were selected and exported for analysis using a custom-built scoring algorithm as described in Hinz et al., 2024.

### Fluorescent and luminescence assays

NAD^+^/NADH and ATP levels were measured using the Promega NAD/NADH-Glo (G9072) and CellTiterGlo (G7572) kits following manufacturer’s recommendations, respectively. Dead cell puncta were measured by adding Incucyte Cytotox Red reagent (4632) to the cell culture medium at a final dilution of 1:4000, capturing both phase and red fluorescence images using the S3 Incucyte Live-Cell Analysis System every 2hrs over a period of 72 hrs and quantifying red puncta using the built-in analysis modules. To measure NfL levels released by cortical neurons into the media, supernatant from neurons was used in NF-L ELISA (Uman Diagnostics, 10-7002) according to the manufacturer’s instructions.

### Animals

All experiments involving animals were approved by Genentech’s Institutional Animal Care and Use Committee. 9-10 week old male and female mice (C57bl/6J) were obtained from Jackson Laboratories. *Sarm1*^-/-^ mice (Kim et al., 2007) were originally obtained from Jackson labs (B6.129X1-*Sarm1^tm1Aidi^*/J) and were maintained on a C57BL/6J genetic background at Genentech. Heterozygous pairs generated *Sarm1*^-/-^ and *Sarm1*^+/+^ animals. Animals were housed on a 14h light/ 10hr dark cycle with *ad libitum* access to food and water.

### Drugs, dosing, and pharmacokinetic analysis

All compounds were administered by oral gavage in a volume of 10 ml/kg. Vacor was administered at 50 mg/kg or 30 mg/kg in vehicle 0.5% methylcellulose / 0.2% Tween 80 (MCT) as established in Loreto et al., 2021. NB-3 was prepared in vehicle 2.5% DMSO, 10% Cremophor EL, and 85% sterile filtered pH 7.0 water with 5% dextrose as described in Batkowski et. al. GNE-5152 was prepared in vehicle 50% Phosal 50 PG, 45% PEG 400, 5% Ethanol. SARM1 inhibitor compounds were dosed 1 hour prior to vacor dosing or SNT surgery, and again 8hrs later. Blood samples for plasma NfL and PK were collected 24hrs following vacor dosing or SNT by retro-orbital bleed into K_2_EDTA-containing tubes. Samples were centrifuged at 17,000 x *g* for 3 min and the supernatant was collected and frozen at −80°C until bioanalysis. Plasma concentrations of compounds were determined using LC MS/MS.

### Vacor model

A single dose of vacor was administered orally and blood samples were collected 24hrs later by retro-orbital bleed. In studies with inhibitor compounds, BEIs were administered 1hr prior to vacor and again 8hrs later. Mice were monitored closely throughout the study period and euthanized immediately if adverse events, such as lethargy, gait disruption, hunched posture, or immobility were observed. Vacor dosed alone at 40 mg/kg and below caused no adverse effects, while 50 mg/kg vacor alone caused adverse effects rarely (∼5% of all animals).

### Sciatic Nerve Transection

SNT was performed based on Bráz et al., 2011. Animals were anesthetized in an induction chamber under a mix of isoflurane and oxygen, and then transferred to a nose cone for anesthesia maintenance under 2.5% isoflurane. The leg was shaved and aseptically prepared with alcohol and Chloroprep. An incision was made at the mid-thigh level on one side and the muscles overlaying the sciatic nerve were separated by blunt dissection. The sciatic nerve was then cut and a 1-2 mm piece was removed. The muscle was closed with a single suture (6-0, Ethilon polyamide 6, non-absorbable) and the skin incision was closed with sutures and Vetbond glue.

### Pharmacokinetic Analysis

The pharmacokinetics of NB-3 and GNE-5152 were determined in male C57Bl6 mice following oral administration. Three male C57Bl6 mice per dose group, aged 6-9 weeks with body weight of ∼20-30 g were given a dose of 30 mg/kg BID (0 hr & 8 hr) of NB-3 formulated in 2.5% DMSO, 10% Cremophor EL, and 85% sterile filtered pH 7.0 water with 5% dextrose; 5 or 50 mg/kg BID (0 hr and 8 hr) of GNE-5152 formulated in 50% Phosal PG 50, 45% PEG400, 5% Ethanol. Following oral administration, 15 µL blood was collected at 0.5, 1, 3, 8, 8.5, 9, and 24 hour post-dose and diluted with 60 µL of water containing 1.7mg/ml Potassium (K_2_) EDTA. Food and water were available to all animals, ad libitum.

Standard curves were prepared in diluted mouse blood and loperamide was used as the internal standard. The standard working solution at 0.1 mg/mL was prepared by appropriate dilution of the 1.0 mg/mL stock solution with DMSO. The loperamide working solution at 10 ng/mL was prepared by appropriate dilution of the 1.0 mg/mL stock solution with acetonitrile. The diluted blood standards, range from 1.0 to 20,000 ng/mL, were prepared by appropriate dilution of the 0.1 mg/mL standard working solution with control diluted blood matrix. Aliquots of 25 µL of diluted blood samples and standard were transferred to the plate and then 400 µL of acetonitrile with internal standard was added to each well. The samples were vortexed and centrifuged.

Concentrations of NB-3 and GNE-5152 in diluted blood were determined by a non-validated LC-MS/MS method. The concentrations of NB-3 and GNE-5152 were determined by a Waters Acquity UPLC System, Sciex Triple Quad 6500 Plus equipped with a turbo-electrospray interface in positive ionization mode. The aqueous mobile phase (A) was 10 mM HCOONH_4_ in water/ACN (v:v, 95:5) and the organic mobile phase (B) was 10 mM HCOONH_4_ in ACN/water (v:v, 95:5). The gradient for NB-3 started with 10% B, and then increased to 90% B in 1.0 minutes, and maintained at 90% B for another 0.2 minutes; decreased to 10% B within 0.01 minutes and maintained at 10% B for another 0.2 minutes. The gradient for G’5152 started with 10% B, and then increased to 95% B in 1.2 minutes, and maintained at 95 % B for another 0.2 minutes; decreased to 10% B within 0.01 minutes and maintained at 10% B for another 0.2 minutes. The flow rate for both compounds was 0.6 mL/min and the cycle time (injection to injection including instrument delays) was approximately 1.6 minutes. A volume of 2 µL of the final extract was injected onto the analytical Waters ACQUITY UPLC HSS T3 Column (50 × 2.1 mm, 1.8 μm).

Quantitation was carried out using the multiple reactions monitoring transition *m/z* 262.1→ 135 for NB-3, *m/z* 366.2→ 144.1 for GNE-5152 and *m/z* 477.1 → 266.2 for loperamide. The optimized instrument conditions included a source temperature of 450°C, a curtain gas pressure of 40 psi, a nebulizing gas (GS1) pressure of 50 psi, a heating gas (GS2) pressure of 50 psi, and a collision energy (CE) of 14 V for NB-3, 75 V for GNE-5152, and 17 V for loperamide. IonSpray needle voltage was set at 5500 V. LC-MS/MS data were acquired and processed using Analyst Software (v1.7.2). The quantitation of the assay employed a calibration curve, which was constructed through plotting the analyte/internal standard peak area ratios versus the nominal concentration of analyte with a weighted 1/(*x*x)* linear regression. PK parameters were calculated by non-compartmental methods using the extravascular model, Phoenix™ WinNonlin^®^, version 6.4 (Certara USA, Inc., Princeton, NJ).

### Plasma NfL measurement

Plasma NfL was measured by a digital SiMoA HD-X immunoassay (single molecule array, Quanterix Corp, Lexington, MA). NF-Light^TM^ v2 Advantage Kit (104073) of the same lot number was used for each experiment. The assay was performed following manufacturer’s instructions. Plasma samples were run at a 1:40 dilution, except for vacor treated samples, which were run at a 1:200 dilution. Reported values are within the limits of detection established by the lot’s Certificate of Analysis datasheet for each experiment.

## AUTHOR CONTRIBUTIONS

R.L., M.W., C.C., S.H., A.C., B.B., K.M., T.K., L.L., Y.C., A.G., J.P.E., A.D. and F.I.H. designed, performed or analyzed experiments. S.G. and G.B. synthesized reagents. S.G., M.N., J.L., A.A.N., B.C and C.C.H. designed experiments. R.L., A.D. and F.I.H. wrote the manuscript.

## ACKNOLEDGEMENTS

We thank Bing-Yan Zhu, Jessica Grandner, Jun Liang, Matthew Del Bel, Mingshuo Zeng, and Russell Smith for their compound design contributions. We thank the chemistry team members at Wuxi, Shanghai for compound synthesis support. We also thank the Genentech DMPK and Compound Management groups for their technical contributions.

