## Supplementary Figures 1-6 for "SARM1 orthosteric base exchange inhibitors cause subinhibitory SARM1 activation"

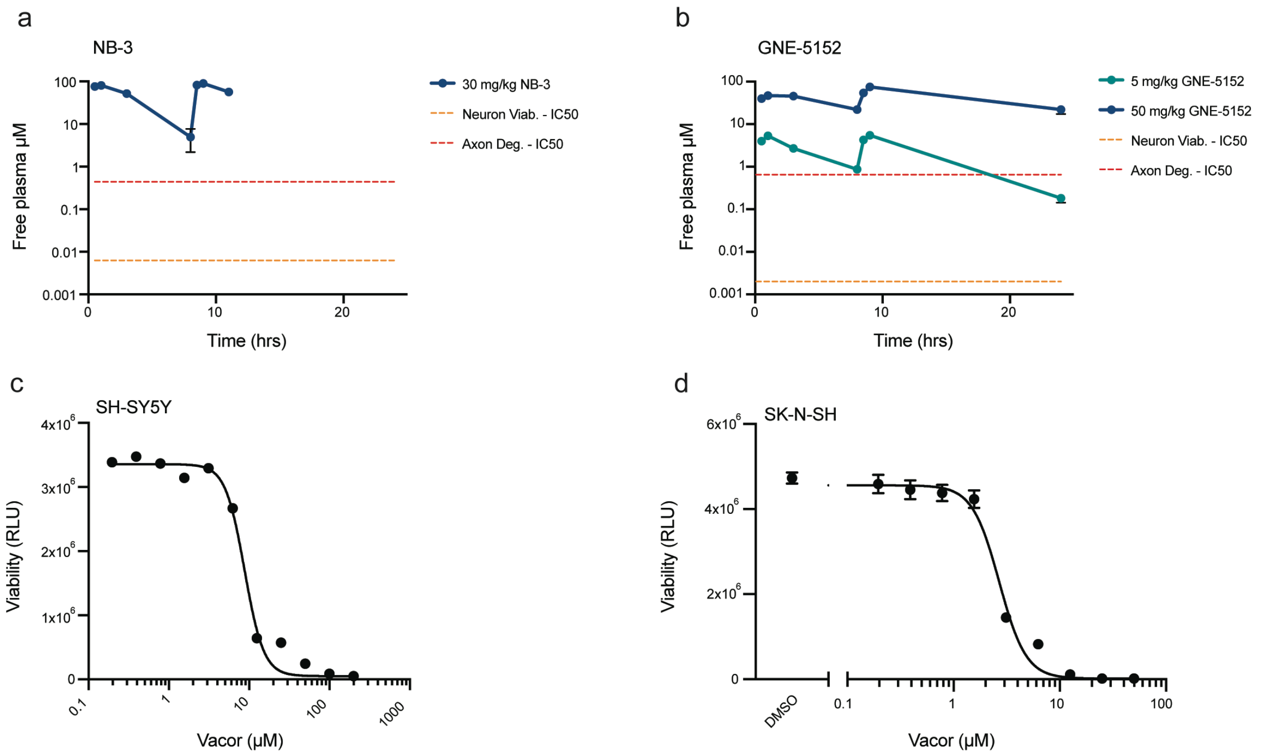


Supplementary Figure 1.

(a) Time course of free plasma concentration of NB-3 after dosing health mice at 30 mg/kg via oral gavage at 0 and 8hrs (measured out to 12hrs). n=3 mice. (b) Time course of free plasma concentration of GNE-5152 after dosing of 5 mg/kg or 50 mg/kg delivered via oral gavage at 0 and 8hrs. n=3 mice. Viability of (c) SH-SY5Y and (d) SK-N-SH cells in response to varying concentrations of vacor.

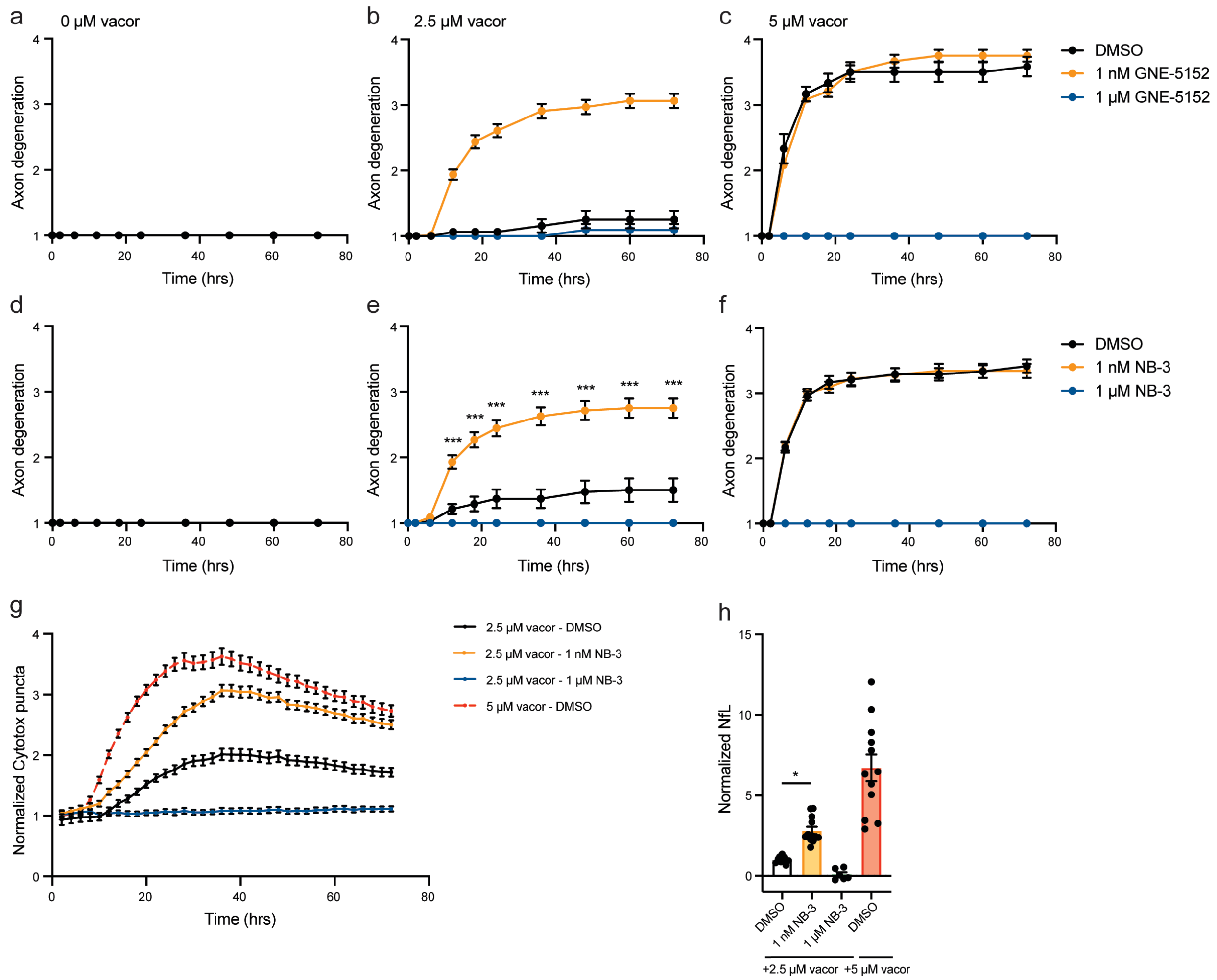


Supplementary Figure 2.

Quantification of axon degeneration of WT ms cortical neurons treated with (a) 0 μM vacor, (b) 2.5 μM vacor and (c) 5 μM vacor in the presence of DMSO, 1 nM or 1 μM GNE-5152. n=12-64 images/condition. Quantification of axon degeneration of WT ms cortical neurons treated with (d) 0 μM vacor, (e) 2.5 μM vacor and (f) 5 μM vacor in the presence of DMSO, 1 nM or 1 μM NB-3. n=16-56 images/condition. (g) Quantification of cytotoxic puncta of WT ms cortical neurons treated with 5 μM vacor or 2.5 μM vacor in the presence of DMSO, 1 nM or 1 μM NB-3, normalized to 0 μM vacor. n= 14-20 images/condition. (h) NfL release from ms cortical cultures 48hrs after treatment, normalized to 2.5 μM vacor + DMSO. n=6-12 wells/condition; Dunnett's multiple comparisons test, *$p<0.05$.

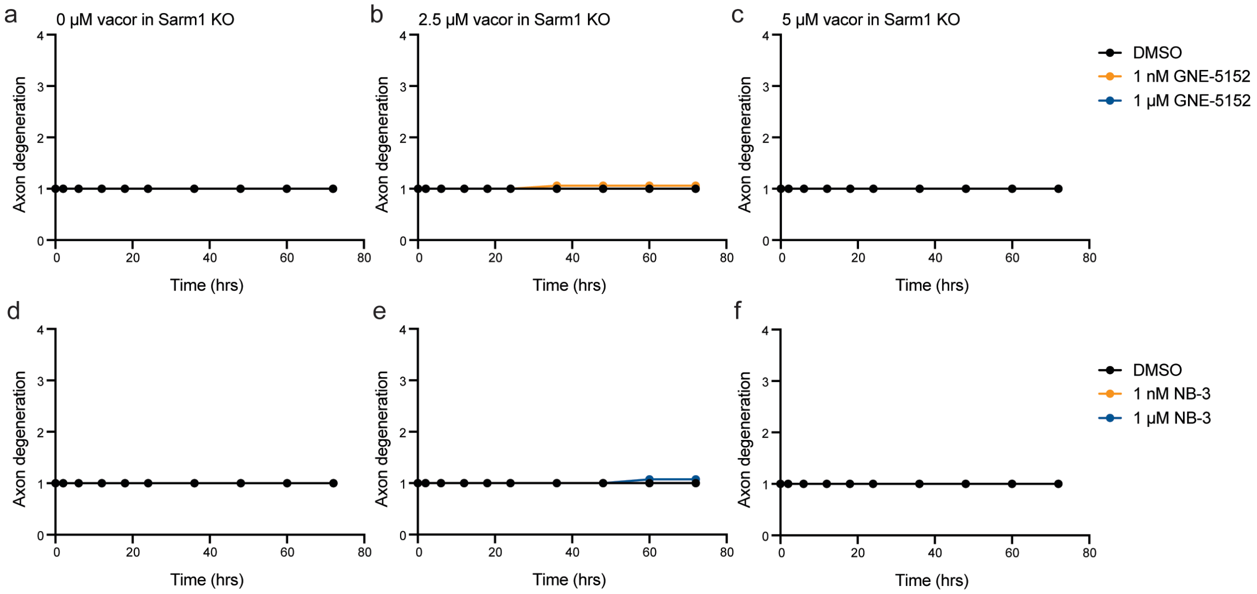


Supplementary Figure 3.

Quantification of axon degeneration of Sarm1 KO ms cortical neurons treated with (a) 0 µM vacor, (b) 2.5 µM vacor and (c) 5 µM vacor in the presence of DMSO, 1 nM or 1 µM GNE-5152. n=22-48 images/condition. Quantification of axon degeneration of Sarm1 KO ms cortical neurons treated with (d) 0 µM vacor, (e) 2.5 µM vacor and (f) 5 µM vacor in the presence of DMSO, 1 nM or 1 µM NB-3. n=18-40 images/condition.

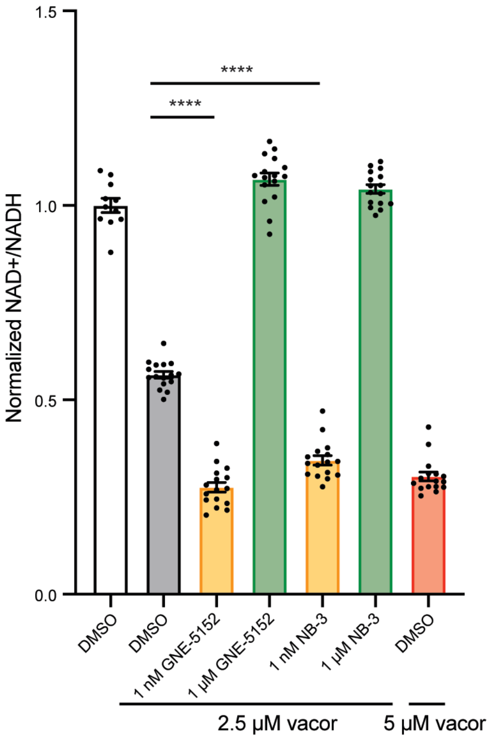


Supplementary Figure 4.

Quantification of NAD+/NADH levels in ms cortical neurons 15hrs after treatment, normalized to DMSO. n=12-16 wells/condition; 2-way ANOVA, ****$p<0.0001$.

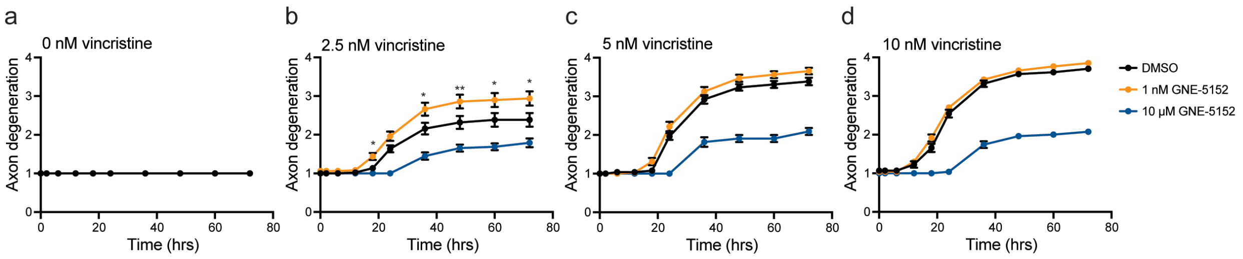


Supplementary Figure 5.

Quantification of axon degeneration of WT ms cortical neurons treated with (a) 0 nM vincristine, (b) 2.5 nM vincristine, (c) 5 nM vincristine and (d) 10 nM vincristine in the presence of DMSO, 1 nM or 10 µM GNE-5152. n=25-50 images/condition; Mann-Whitney test, *p<0.05, **p<0.01 comparing 2.5 nM vincristine + DMSO to 2.5 nM vincristine + 1 nM GNE-5152.

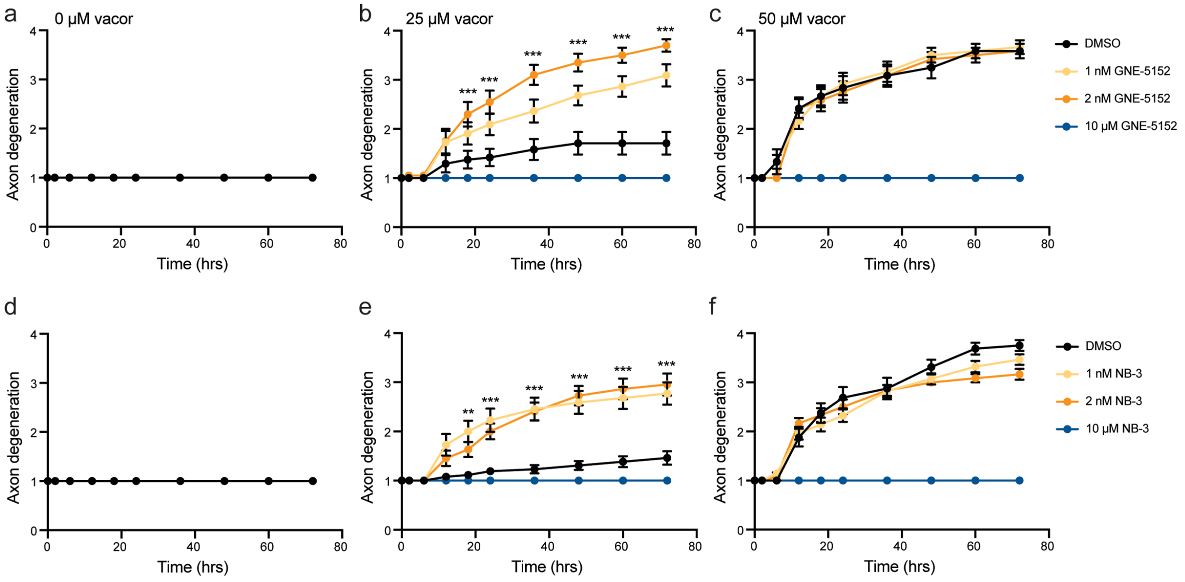


Supplementary Figure 6.

Quantification of axon degeneration of WT human iPSC-derived cortical neurons treated with (a) 0 µM vacor, (b) 25 µM vacor or (c) 50 µM vacor in the presence of DMSO, 1 nM, 2 nM or 1 µM GNE-5152. n=12-32 images/condition. Quantification of axon degeneration of WT human iPSC-derived cortical neurons treated with (d) 0 µM vacor, (e) 25 µM vacor or (f) 50 µM vacor in the presence of DMSO, 1 nM, 2 nM or 1 µM NB-3. n=12-28 images/condition. Mann-Whitney test, *p<0.05, **p<0.01, ***p<0.001.
